# Population-level sensitivity to landscape variables reflects individual-based habitat selection in a woodland bat species

**DOI:** 10.1101/752733

**Authors:** Pierre-Loup Jan, Diane Zarzoso-Lacoste, Damien Fourcy, Alice Baudouin, Olivier Farcy, Josselin Boireau, Pascaline Le Gouar, Sébastien J. Puechmaille, Eric J. Petit

## Abstract

Understanding the relationship between habitat quality and population dynamics is fundamental for long-term management and range predictions in ecology. However, habitat quality is generally only investigated at the individual scale, as it is the case for the lesser horseshoe bat (*Rhinolophus hipposideros*), a species of conservation concern. Using a statistical modelling approach and census data of 94 lesser horseshoe bat colonies located in Brittany (France), we analysed the effect of landscape composition and configuration on the demographic components of surveyed maternity colonies (i.e. colony size, fecundity and growth rate), and compared our result to those provided by individual-based studies. Our results validated that the landscape in a 500-meter buffer around colonies (core foraging area) is crucial for population size and dynamics, and confirmed the positive influence of broadleaved woodland proportion on bat colony size. We revealed a positive effect of lakeshores and riverbanks on colony size and growth rate, underlying the importance of these habitats for the long-term conservation of this non-migratory forest species. Importantly, our results refine previous knowledge concerning the threat posed by the intensification of human activities (e.g. urbanization, agriculture, habitat fragmentation), and highlight the negative effect of large and regular patches of artificial and crop lands and of open land patches shape complexity on all demographic variables investigated. While our results support the dependence of population dynamics and associated conservation management to individual behaviour and sensitivity, environmental responses differed between the population metrics investigated, showing that efficient range prediction will require to fully grasp the complexity of the interaction between landscape and the different population dynamic parameters.

## 1. Introduction

Better understanding the relationship between population dynamics and habitat quality is fundamental both for predicting range shifts under global change and for establishing sound conservation strategies (Elith and Leathwick, 2009; Hodgson et al., 2009). Recently, Species Distribution Modelling (SDMs) has become the standard to inform scenarios of biodiversity under global change (e.g. climate and land use changes) by correlating species occurrence with eco-geographical information related to species climate and/or habitat niche (Elith and Leathwick, 2009). These models, however, do not explicitly consider the demographic processes (e.g. reproduction or survival) relating population-level metrics (e.g. presence of a population, density, population growth rate, carrying capacity) to habitat quality (Thuiller et al., 2013), while they are essential for species to follow environmental conditions favourable to their survival (species range dynamics-Aitken et al., 2008; Auffret et al., 2017; Gaillard et al., 2010; Kokko and López-Sepulcre, 2006). Therefore, research questions that target population-level processes should use metrics that relate to population dynamic parameters (Gaillard et al., 2010).

Yet, studies that directly evaluate relationships between population-level metrics and habitat quality are seldom both because demographic data are not often gathered over many populations (Gurevitch et al., 2016; Bergman et al. 2019) and because landscape variation is still difficult to assess at a scale fine enough to meaningfully represent the diversity in habitat quality (Anderson 2018). If population-level species-habitat relationships are so rarely documented, where does our basic ecological knowledge come from? Historically, from the study of resource selection by individuals (Manly et al., 2007).

The fact that population-level studies of habitat quality are not frequent has triggered evaluations of the necessary conditions for using one metric (i.e. population-level metrics) as a proxy for another (i.e. habitat quality - Thuiller et al., 2014). For instance, though neither species occurrence neither resource selection by individual directly relates population-level dynamics to environmental conditions, both have been used to predict abundance (Acevedo et al., 2017; Boyce et al., 2016; Street et al., 2016). Studies relating population-level metrics and processes to habitat quality are needed to test their validity as surrogates for inferring this relationship. More precisely, scaling up resource selection by individuals to abundances is informative, at least when environmental variables that are important for variations in population sizes were included in the study of resource use (Boyce et al., 2016; Street et al., 2016). In general, establishing in which conditions environmental variables that come out of individual-based studies are also relevant at the population-scale would give more weight to conservation actions based on data gathered at the individual level, and enlarge the range of studies that can feed more realistic predictions of species range shifts.

The lesser horseshoe bat, *Rhinolophus hipposideros,* has undergone a massive decline in the northern part of its distribution range during the last century for reasons that are still not fully understood, although pesticides use, decline in insect prey, and habitat destruction were suspected causes (Bontadina et al., 2000). This stressed the need for a better understanding of its habitat requirement, leading to numerous telemetry studies (e.g Bontadina et al., 2002; Downs et al., 2016; Reiter et al., 2013; Schofield, 1996) and, more recently, to larger scale presence/absence studies (Le Roux et al., 2017; Tournant et al., 2013). These studies mainly focused on habitat selection by females during the crucial period of parturition, during which they gather in maternity colonies. Individual-level studies revealed that the lesser horseshoe bat forages in woodland, especially broadleaved ones, and spend most of their foraging time in close vicinity to their roost (within a radius of 500-600 meters - Bontadina et al., 2002; Reiter et al., 2013). The importance of woodland proportion on the presence and abundance of *R. hipposideros* was confirmed by several studies (e.g. Reiter, 2004; Tournant et al., 2013), and presence data revealed a negative effect of artificial land cover on the presence of maternity colonies (Tournant et al., 2013). Habitat requirements of this species may dramatically constrain the opportunity offered by a changing climate to its predicted northward expansion (Rebelo et al., 2010).

In this study, we tested the hypothesis that important environmental variables known from individual radio-tracking studies or presence surveys also influence the demography of the species. We used count data obtained from a fifteen years monitoring scheme at 94 colonies situated in Brittany (France), to investigate the population-habitat relationship in *R. hipposideros*. We used these data to infer variables that are related to the population dynamics of the colonies before testing their relationship with environmental variables.

## 2. Material and Methods

### 2.1. Demographic data for *R. hipposideros* colonies

From 2000 to 2014, 94 *R. hipposideros* maternity colonies were monitored in Brittany, France (Fig. 1). Monitoring were performed by volunteers from two NGO, Bretagne Vivante and Le Groupe Mammalogique Breton, who followed the same protocol for all colonies and performed visual counts with minimum disturbance during late June or early July, that is, during the period when new-borns are easily distinguished from adults in Brittany. One or two counts could be performed each year, depending on colony accessibility and volunteer availability, and when multiple counts were carried out in a given year, only the largest one was considered. Adults and juveniles were counted separately: each year, the census size of the colony was estimated by the number of adults, and the fecundity by the number of juveniles divided by the number of adults. Genetic sexing on *R. hipposideros* has shown that while adult male can be detected in colonies, the duration of their presence are so short that adult visual counts can be considered as a good estimate of the number of adult females (Zarzoso-Lacoste et al., 2018). Not every colony was known in 2000 and, in some cases, monitoring was not possible due to unforeseen circumstances (blocked access to the bats or the person in charge of counting). Thus, the number of monitored years per colony ranged from 3 to 14 (7.73 on average). The adult females that compose these colonies are highly philopatric (Dool et al., 2016), and dispersal between maternity colonies is rare enough that we can consider them as closed entities (Jan et al., 2019). Thus, we can consider maternity colonies as different populations, in a sense that they consist of groups of individuals of the same species that co-occur in space and time, have an opportunity to interact with each other, and sufficiently isolated that immigration does not substantially affect the population dynamics (Waples and Gaggiotti, 2006). Temporal variation of colony size and fecundity in this dataset, as well as their dependency on environmental covariates, were already investigated in Jan et al. (2017). In this paper, we focused on inter-colony variation instead and their dependency to environmental covariates that were constant over time, the landscape structure and configuration. Hence, the following demographic variables were compiled for each colony: mean colony size (the number of adults averaged over years), fecundity (the juveniles/adults ratio averaged over years), and growth rate. The last variable was estimated with a Malthusian model using the package “PVAclone” (Nadeem and Solymos, 2016), which allows 1) to compute the maximum likelihood estimates and corresponding standard error using a cloning algorithm and 2) to use a general state-space model formulation to assume the presence of observation error while fitting the model. In our computation, five clones were considered in the data-cloning algorithm, 20 000 iterations were performed, and the models were fitted assuming that the observation error followed a Poisson distribution. The Malthusian model is density-independent and considers that the observed colony size variation through time is only the result of growth rate and observation error, which allows to estimate the growth with a known precision. The reasons behind the choice of the model are detailed in Supp. Inf. A.

**Figure 1:**
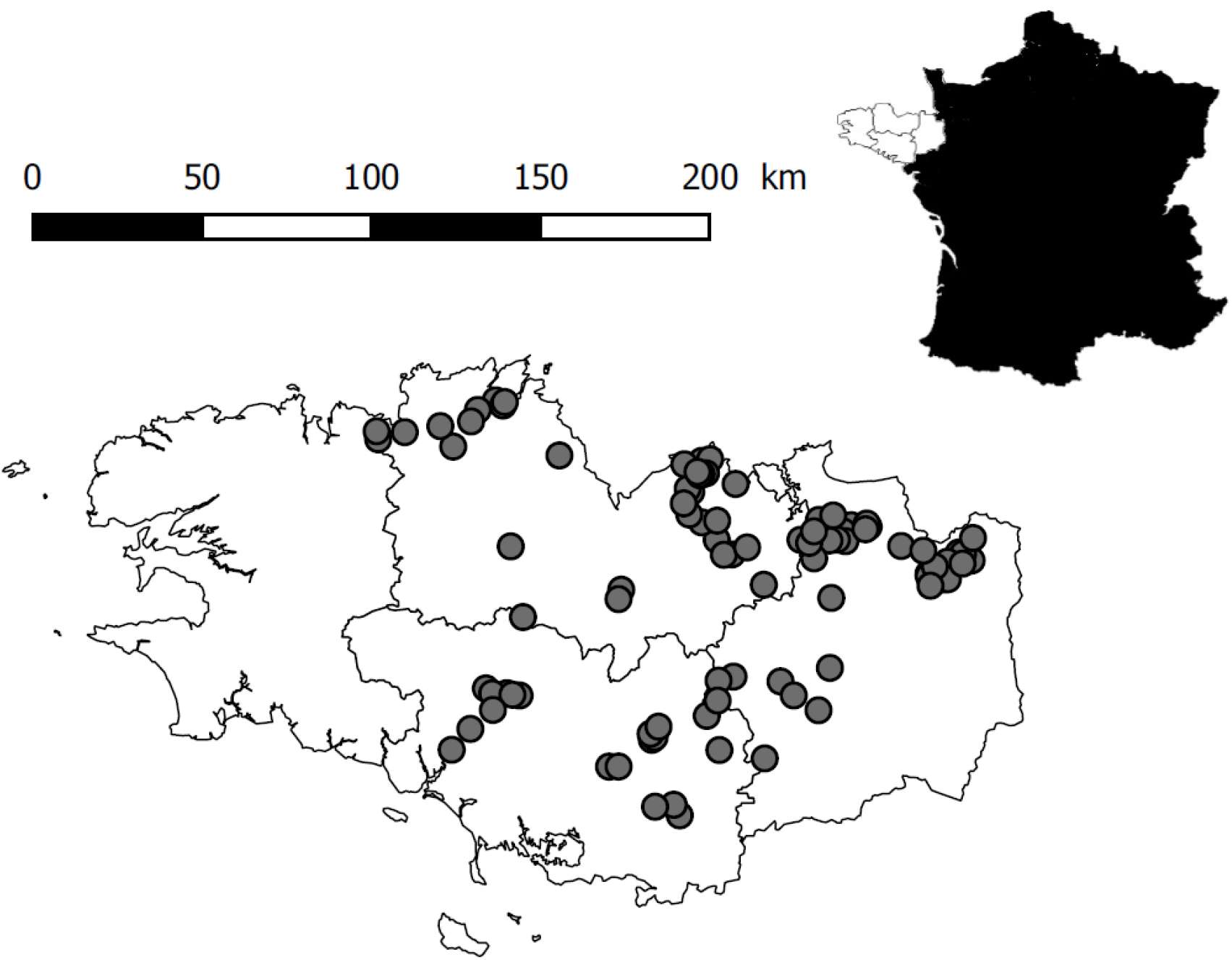
Map of the 94 colonies monitored in Brittany, France.

### 2.2. Landscape attributes

We built a geographic information system (GIS) of land cover associated with other descriptors of the landscape surrounding the colonies. To obtain information precise enough and relevant for *R. hipposideros,* the land cover dataset was built from different databases: CORINE Land Cover - European Commission 2006, BD Topo 2.1 - IGN 2013, BD Forest v1 - IGN 2012 and Graphic Parcel Register 1.0 (RPG) - IGN 2012. The land cover map construction included procedures for correcting topological errors, homogenizing information resolution and harmonizing attribute data. Finally, we obtained a map featuring 38 classes of land use (Supp. Inf. B) with a minimum mapping unit (MMU) of 100 m^2^.

### 2.3. Assessment of the effect of landscape composition and configuration on bat population dynamics

We considered the landscape within a circular buffer surrounding each colony. Two buffer radius were tested: 2500 m, reflecting the radius commonly used in recent studies on the lesser horseshoe bat (Afonso et al., 2016), and 500 m, reflecting the core area of the foraging zone (Bontadina et al., 2002; Reiter et al., 2013). Before clipping the land cover layer with each buffer, the 38 cover types were aggregated in six land-cover classes (Supp. Inf. B) according to previous knowledge gathered at the individual level on lesser horseshoe bat’s ecology (Bontadina et al., 2002; Tournant et al., 2013), namely: broadleaved woodland, coniferous woodland, artificial area, water bodies, cropland, and open land (other than crops and artificial). For each buffer we calculated six FRAGSTATS metrics (McGarigal and Marks, 1995) describing the composition and the configuration of the landscape: the proportion of the landscape occupied by each class (%LAND), the mean patch size (MPS) and the mean SHAPE index (MSI) of each class, and three landscape diversity metrics, namely: the Simpson’s diversity index (SIDI), the Shannon’s diversity index (SHDI) and the Shannon’s evenness index (SHEI). Mean patch size was calculated by averaging the surfaces of all homogeneous areas (patches) of all land-cover classes present in the buffer. Because woodland and riparian vegetation are highly selected habitats, while artificial, open and crop land are avoided, as documented in individual-level studies (Bontadina et al., 2002; Tournant et al., 2013), we hypothesised that the surface of these land-cover classes is positively (for selected surfaces) or negatively (for avoided ones) related to population-level metrics. Habitat selection studied in another European woodland specialist bat has shown that woodland exclusively composed of conifers could be avoided instead of selected (Smith & Racey 2008). As a consequence, we considered coniferous woodland separately in our analysis. MPS and indices of landscape diversity are easy-to-compute generalist landscape configuration metrics, not dependent on land classes, widely used in landscape ecology studies. Low MPS is supposed to reflect general landscape fragmentation (Wang et al. 2014), while higher landscape diversity, by implying more edge effect in the landscape, has already been associated with an increase in *Rhinolophidae* activity (Ancillotto et al. 2017). Since a MSI value of 1 corresponds to a circular patch, and because this value increases with patch shape irregularity, a high MSI value reflects an increase in edge density of a particular land cover, for an equivalent area. MSI of selected habitat would measure the edge effect (which is expected to increase the foraging opportunity of insectivorous woodland bat - Morris et al. 2010) or the effect of particular configuration prone to affect woodland bat abundance, while those of avoided habitat can be considered as a proxy for habitat fragmentation (Wang et al. 2014).

Statistical model averaging was then performed to estimate the effect of land cover variables on population dynamic parameters. Full models considered either the colony size (Generalized linear model with negative-binomial distribution - Jan et al. 2017), fecundity (linear model) or the growth rate (weighted least squares regressions) as the response variable, and the proportion and mean shape index of the six land cover, mean patch size and land cover diversity as explanatory variables (without interaction). Explanatory variables were centred and scaled before model computation. Weighted least square regressions were performed to take into account the uncertainty around growth rate estimates by weighting each data point by the inverse of the standard deviation given by the estimation (Ryan, 2008).

These models were then used in the glmulti R package to obtain every possible combination of explanatory variables (without interaction) and order them by AICc criterion (Calcagno and de Mazancourt, 2010). We then performed model averaging by calculating the Akaike weight of each model within 2ΔAICc of the best model (Burnham and Anderson, 2003). The model-averaged regression coefficients of the predictors and their 85% confidence intervals (CI) were calculated based on the cumulative weights of the models including the variable (Calcagno and de Mazancourt, 2010). 85% confidence intervals are more relevant than 95% interval confidence for AIC-based model averaging (Arnold, 2010), and explanatory variables were then considered as having a meaningful positive or negative impact on the response variable if their 85% confidence interval did not include zero. Because growth rates and their associated standard deviations obtained through PVAClone can slightly vary from one run to another, we decided to run ten times the growth rates estimation and the model averaging associated with this response variable. Only the explanatory variables that were significant ten times out of ten were considered as significant in our results.

Land cover areas were considered through their proportions, inevitably leading to correlation between variables. We prevented this by discarding models including more than four proportion variables for model averaging. Correlations between coefficients were lower than 0.6 for every remaining models. The three diversity indexes (SIDI, SHDI and SHEI) were also highly correlated, and we selected the index which explained most variability for each demographic variable by building simple linear models with every response variable prior to model averaging.

Since we previously showed that some climatic variables impact colony size and fecundity (Jan et al., 2017), we averaged these variables (June and October precipitation and May and November mean temperature for Colony size; April and October precipitation and April and July minimum temperature for fecundity) along the sampling period for each colony and included them in all models using colony size or fecundity as a response variable to control for the variability explained by the weather. The description of the complete models which were the basis of model averaging are presented in Supp. Inf. C. Our data set provides population dynamic parameters on a large spatio-temporal scale, and a sample size large enough to investigate numerous environmental variables. Complete models included the whole 94 colony dataset to test the significance of no more than 14 environmental variables (climatic variables being present in the model but not tested), ensuring than the ratio sample size (*N*) / tested predictor (k) was high enough to avoid overparametrisation (N/k = 6.71, Forstmeier and Schielzeth, 2011).

To determine which radius (500 or 2500m) best explained variations in our response variables, we compared the AICc and R^2^ of the models explaining the population dynamics parameters with the significant predictors detected in the previous model averaging.

## 3. Results

Shannon’s evenness index (SHEI) was the diversity index explaining most variability for the three demographic variables, and was thus the only diversity index included in model averaging. Significant landscape variables were detected for each response variable when considering a 500 m radius buffer around the colony: four for colony size, one for fecundity and six for the growth rate (Tab. 1). The proportion of artificial land negatively impacted the three investigated response variables. The growth rate was also negatively impacted by the proportion of crop areas, while colony size was positively influenced by the proportion of broadleaved woodland. Mean Shape Indices (MSI) associated with water bodies were positively related to colony size and growth rates, which were also positively influenced by the MSI of crop and artificial lands. Finally, MSI of open land had a negative impact on colony size and growth rates.

**Table 1:** Results of a model averaging of population dynamics parameters as a function of the proportion of land (%LAND) and Mean Shape Index (MSI) of different cover type, the mean patch size and SHEI (Shannon’s evenness index). NS: non-significant predictor; ‘+’: positive significant predictor; ‘-’: negative significant predictor. When results differ between the two buffer sizes, results given by the 2500m buffer are indicated between brackets. Cells with significant results with the 500m buffer are shaded.

|  | Broadleaved woodland |  | Coniferous woodland |  | Artificial |  | Crop land |  | Open land |  | Water bodies |  | SHEI | Mean patch size |
| --- | --- | --- | --- | --- | --- | --- | --- | --- | --- | --- | --- | --- | --- | --- |
|  | %LAND | MSI | %LAND | MSI | %LAND | MSI | %LAND | MSI | %LAND | MSI | %LAND | MSI |  |  |
| Colony size | + | NS | NS | NS | - (NS) | NS | NS | NS | NS | - (NS) | NS | + (NS) | NS | NS |
| Fecundity | NS | NS | NS | NS | - (NS) | NS | NS | NS | NS | NS | NS | NS | NS | NS |
| Growth rate | NS | NS(+) | NS | NS | - (NS) | + (NS) | - (NS) | + | NS | - (NS) | NS (+) | + (-) | NS | NS (+) |

Only six significant predictors were detected within the 2500 m radius buffer, two of them being concordant with the results of the 500 m radius buffer (Tab. 1). When comparing the AICc and R^2^ of the model given by the two buffer sizes, fixed effect detected with the 500m buffer always performed better (Tab. 2). Effect size plots are presented in Suppl. Inf. D. For the 500 m radius buffer, models constructed with significant land cover predictors explained 42% of the variance for colony size, and 8% for fecundity. These values reached 53% and 13% when significant climatic predictors previously detected for colony size and fecundity were added to the corresponding models (Jan et al., 2017). The six significant predictors found in growth rate models explained 80% (±3%) of this variable, with crop land proportion being the predictor with the largest effect size and explaining 53% (±5%) of the variation.

**Table 2:**
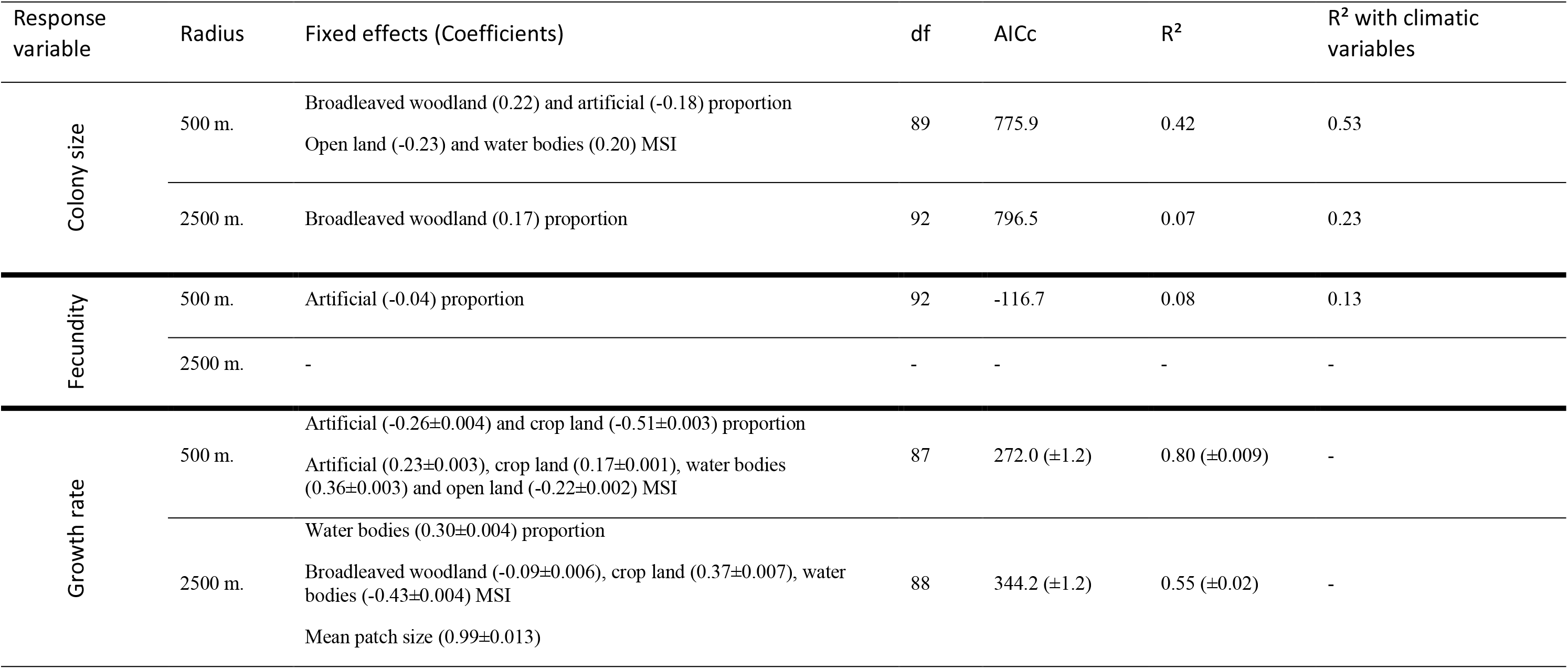
Coefficients, Residual degree of freedom (df), AICc and R^2^ of models that included land cover significant predictors for the two buffer scales. For colony size and fecundity, R^2^ of models that include significant climatic predictor detected in a previous study (Jan et al. 2017) are also presented. Coefficients, AICc and R^2^ of growth rate model are averaged over the ten runs we performed for growth rate calculation, and presented with the corresponding standard error (See Material and Methods).

## 4. Discussion

### 4.1. Scale of habitat dependency and management implication

We showed that investigating the effect of landscape composition and configuration on the dynamics of *R. hipposideros* maternity colonies is more relevant at the smallest of the two investigated scales. Indeed, the AICc and R^2^ values of models including landscape significant predictors always supported the importance of the landscape surrounding colonies within a 500 m radius (lower AICc and larger part of explained variance R^2^ - Tab. 2). This result is consistent with those of individual level telemetry studies demonstrating that individuals forage for at least half of their time within a 500-600 m radius around maternity roosts, although few individuals can eventually forage at distances up to 5 km around colonies (Bontadina et al., 2002; Reiter et al., 2013). Nevertheless, further investigations are required to precise the most relevant spatial scale at which habitat (affecting colony sizes and dynamics) should be protected or managed for an optimal and cost effective conservation of this species, while taking into account connectivity between populations (Lehnen et al., 2021).

### 4.2. Identification of key habitats for *R. hipposideros* population dynamics and conservation

Colony size was positively influenced by broadleaved woodland proportion at both spatial scales. This validates and refines previous studies, in which woodland areas was one of the best landscape predictors of maternity roost presence (Bontadina et al., 2002; Reiter, 2004; Schofield, 1996; Tournant et al., 2013). Indeed, broadleaf woodlands, often offering a higher richness and abundance of insects than coniferous plantations (Benton et al., 2002; Goiti et al., 2004), are more intensively used as foraging areas by *R. hipposideros* than any other habitat (Bontadina et al., 2002; Schofield, 1996).

We also found a significant positive effect of water bodies shape complexity on colony size and growth rate. Because our GIS computation lead to a minimum mapping unit (MMU) of 100 m^2^, small rivers and ponds are not considered in our models, and distinction between standing and flowing water is not possible. This means that the positive effect detected reveals that long lakeshores, riverbanks, or both of them near maternity colonies favour colony size and the long-term establishment of this species. Shores with sufficient riparian vegetation to provide food and shelter to a great density of insect is known to drive the distribution and foraging behaviour of insectivorous bats (Warren et al., 2000), as observed in *R. hipposideros* (Bontadina et al., 2002; Holzhaider et al., 2002; Reiter, 2004). These habitats must represent high quality foraging areas for this bat since a non-negligible part of its main prey (e.g. Trichoptera and Nematocera – Arlettaz et al., 2000; Lino et al., 2014) have an aquatic larval development.

The preservation of large broadleaved woodland and high density of lakeshores and riverbanks in the direct vicinity of maternity colonies appears to be the priority for conserving viable colonies of this non-migratory forest species. These high quality cover types, in terms of resources and protection, are suspected to positively influence bat population dynamics by enhancing individual survival. This knowledge would be highly beneficial to objectively guide bat conservation, for instance by identifying where to favour the settlement of new colonies through roost construction (e.g. Mering and Chambers, 2014) and/or by prioritising the management or renovation of roosting sites (buildings, abandoned mines, caves - e.g. Lacki, 2000).

### 4.3. Landscape variables threatening *R. hipposideros* population dynamics and conservation

Our study highlights that fragmentation and degradation of habitat are a major concern for *R. hipposideros* dynamics.

Indeed, our results revealed a significant and negative effect of open land MSI on population size and growth rate. This means that the complexity of the spatial configuration of open land patches (e.g. patch number, shape) surrounding maternity colonies, as a measurement of habitat fragmentation (Wang et al., 2014), affects the dynamics and persistence of this species. This may reveal a barrier effect of this generally avoided cover type (Bontadina et al., 2002; Holzhaider et al., 2002; Reiter, 2004; Tournant et al., 2013), possibly by hindering individuals’ foraging movements, especially if forest corridors and hedge rows are scarce (Downs and Racey, 2006; Fischer et al., 2010; Lumsden and Bennett, 2005). Indeed, most insectivorous bats are known to use scattered trees or hedgerows as transit and foraging corridors because these natural vertical structures offer shelter from predators and generally present higher insect densities than open areas (Downs and Racey, 2006; Fischer et al., 2010; Lumsden and Bennett, 2005). This seems to be supported by the apparent inability of this species to adapt its spatial foraging behaviour in degraded landscapes, and by the high energetic cost of flight that probably constrains its travelling distance, especially for pregnant or lactating females that mostly compose maternity colonies (Rainho and Palmeirim, 2011; Reiter et al., 2013).

Impact of habitat quality degradation could be assessed with the effects of artificial and crop lands. Proportions of both cover types have negative impact on *R. hipposideros* dynamics. In addition, their complexity indexes have positive effect which suggests a negative impact of large and regular urban or crop patches around the colonies probably by decreasing the edge density and the interactions with other more favourable types of landscape cover. The negative effect of urbanization on colony sizes, fecundity and growth rates, is in accordance with a study based on occupancy data only (Tournant et al., 2013). It may be due to the low quality of this cover type (scarcity of favourable foraging areas, low quality or quantity of food resource, higher exposition to light pollution, accidental mortality, predation, disturbance, stress or pollution – see Boyes et al., 2021; Fensome and Mathews, 2016; Gaston et al., 2013; O’shea et al., 2016; Stone et al., 2009; Ziembicki et al., 2015).

The major and negative effect of crop land proportion on maternity colony growth rates that we detected is probably due to the unfavourable nature of this habitat for *R. hipposideros* (Bontadina et al., 2002). Direct and indirect effects of agriculture intensification have been proposed as one of the main cause implicated in the decline of various organisms (Bayat et al., 2014; Gibbons et al., 2014; Hallmann et al., 2014) including *R. hipposideros* (Bontadina et al., 2000; Wickramasinghe et al., 2004). Pesticides can have various additive or synergistic deleterious effects on survival and health of long-lived insectivorous bats (Hsiao et al., 2016). Their use in conventional farmlands reduces insect abundance and diversity (especially three key families for many insectivorous bats: lepidopteran, dipteran, coleopteran) compared to organic farmlands, the latter being more frequently used as foraging sites by *R. hipposideros* (Wickramasinghe et al., 2004). Agrochemicals also impact insectivorous bats by direct poisoning through dermal contact, inhalation or ingestion in their diet, and because of the accumulation and biomagnification of these pollutants in bat organism, and their remobilization from fat tissues during hibernation or migration (Bayat et al., 2014; Mineau and Callaghan, 2018). During lactation, juveniles strongly accumulate organochlorine pesticides (Lüftl et al., 2005; Streit et al., 1995) through the feeding from their mother, and their survival is more affected by pesticide contamination than adult survival (Frick et al., 2007). Moreover, when foraging, inexperienced juveniles fly at much greater distances and are less selective on habitat quality than reproductive females (Bontadina et al., 2002), and are therefore potentially more exposed to factors affecting their survival (e.g. low prey availability, agrochemicals, predation) in such unfavourable habitats. Previous work on *R. hipposideros* maternity colonies population dynamic has shown that juvenile survival variation was the main driver of growth rate variation (Jan et al. 2019), and there is no doubt that an environment prone to reduce juvenile survival would in turn negatively affect population growth rates.

### 4.4. Consistency between individual- and population-level effects of habitat quality

Understanding how landscape elements influence the dynamics of maternity colonies is a key information both for the long-term conservation of this non-migratory species and to refine range shift predictions under climate change. We observed a positive relationship between habitat quality and bat abundance/density that was consistent with predictions made from individual-based studies: habitats in close vicinity to the colony and predicted as high quality by individual-level studies (i.e. broadleaved woodlands and long lakeshores or riverbanks) were associated to large colony sizes, contrarily to artificial lands, predicted as low quality habitats and associated with small colony sizes, but also with low fecundity and/or negative growth rate. Our results thus support conservation guidelines drawn from individual-based studies by showing that such guidelines (e.g. Reiter et al. 2013) should result in healthy colonies, from a population dynamic point of view.

Habitats area and distribution are however undergoing global changes. This has prompted the development of anticipatory predictive models (*sensu* Mouquet et al. 2015), which use species occurrence or habitat selection to forecast changes in species distributions. The consistency of the relationship between population-, on the one side, and individual-level metrics, on the other side, with environmental variables justify the use of anticipatory models built from occurrences or habitat selection, in species such as *R. hipposideros*. In this species, adding fine scale habitat to climatic requirements would certainly help improve current models (Rebelo et al. 2010).

However, recent studies on species distribution models suggest that performing proper prediction will certainly require to go beyond the simple association between expected landscape changes and habitat selection by individuals (Franklin 2010, Mieszkowska et al. 2013). Habitat requirements are not the same for the dispersal, establishment or long-term persistence of populations. Therefore, developing more accurate species distribution model require to grasp the complexity of the interaction between landscape and the different population dynamic parameters (Franklin 2010).

Interestingly, we observed that the different population metrics we investigated were not equally sensitive to variation in landscape variables, highlighting the need to use several and complementary population dynamics metrics. Broadleaved woodland, which has been demonstrated in numerous individual-level studies as being by far the habitat with the highest quality for *R. hipposideros* foraging and the best indicator of maternity colony presence, was only significantly impacting colony size and not the population dynamic metrics, fecundity and growth rate. Cropland proportion was not found to significantly explain variation in colony size and fecundity despite the strong relation it had with growth rate, which is consistent with our former hypothesis about the impact of cropland area and associated pesticides with survival, and particularly juvenile survival (Jan et al. 2019). The faster temporal turnover of agricultural as compared to forest patches could result in colonies becoming ecological traps when large colonies (linked to broadleaved forests) produce juveniles with low survival probability (because surrounding agricultural patches were converted, for instance, from grassland to cropland).

Indeed, nonequilibrium situations, such as the ecological trap described above, transient dynamics (expanding or contracting populations), or source-sink dynamics, can all blur the relationships between habitat selection, abundance and population dynamics (Boyce et al., 2016). Each population has its own history and faces different challenges, which can cause great variation in their dynamics. Our dataset is no exception, and every colony we included in our analysis did not follow the same dynamics (Supp. Inf. A): some were stable, some were exponentially growing or declining, and others exhibited a more complex pattern (logistic growth). Such diversity is obviously unavoidable when analysing large datasets, but should raise caution about the interpretation of our results. To investigate how this variation could affect our results, we included the different type of population dynamics (stable population, Maltusian growth and logistic growth) as a random effect in our colony size and fecundity analysis models (Supp. Inf. E). These additional results were qualitatively equivalent to the results of the models presented in the Result section. We found exactly the same covariates to be significant, for both scales. While we cannot completely exclude the possibility of missing some environmental effect because of the noise caused by the diversity of population dynamic patterns, this second analysis increases our confidence in the validity of our results.

We know that habitat selection can affect individual fitness and population dynamics through very diverse demographic processes (e.g. sex or stage specific survival rates, fecundity - Bjørneraas et al., 2012; Dyson et al., 2018; Jacques et al., 2015; King et al., 2006; Low et al., 2010; Street et al., 2016). Considering the variety of the possible mechanisms, mediated by individual fitness, driving population-level response to habitat quality, we strongly encourage further cross-validation studies aiming at testing these relationships, for example using a habitats-to-populations (HTP - Matthiopoulos et al., 2019) or Environment-Physiology-Demography (EPD – Bergman et al., 2019) framework. This will help better understand the ecology and population dynamics of the species for which we aim to predict their fate from current knowledge.

### 4.5. Conclusion

We underlined the importance of broadleaved woodland proportion and lakeshores and riverbanks density in *R. hipposideros* core foraging area (500 m radius around maternity colonies), probably because their associated vegetation provides particularly favourable shelter, movement and foraging corridors. We also pinpointed the threat posed by the intensification of human activities (i.e. urbanization and agriculture intensification inducing habitat fragmentation), resulting in the spread of unfavourable and degraded habitats (i.e. with low availability of their prey, high exposure to predation and pollution), on the long-term persistence of large *R. hipposideros* populations.

The links we found between landscape variables and the demography of this non-migratory forest species will help refining range shift predictions based on climatic variables only (Rebelo et al., 2010). Concomitantly, considering the connectivity of favorable habitats and the threats posed by the intensification of human activities (urbanization, agriculture, pollution), for example under different land use change scenarii, will allow to more accurately predict the fate of the species towards the end of the century.

## Acknowledgements

We are very grateful to all volunteers from Bretagne Vivante and Le Groupe Mammalogique Breton for providing visual count data of maternity colonies in Brittany that allowed us to perform this study as well as the companion study published in Oecologia (Jan et al. 2017). We thank Benoît Persyn and Météo-France for providing us with meteorological data through the research convention Météo-France/INRA. This work benefited from funding by the French National Forest Office (ONF). We also acknowledge Brock Fenton, Jean-Michel Gaillard and an anonymous PCI reviewer for useful comments that helped clarify and strengthen the manuscript.

## Data accessibility

R scripts used in this study were deposited at Figshare through the DOI 10.6084/m9.figshare.8337437

## Supplementary Information

Supplementary Information A: Selection of population growth model.

Supplementary Information B: Aggregation of 38 cover types in six-landscape class.

Supplementary Information C: Description of the complete models used for model averaging.

Supplementary Information D: Effect size plots for the three best models investigating the effect of landscape on colony size, fecundity, and growth rate in *Rhinolophus hipposideros* maternity colonies.

Supplementary Information E: Analysis of colony size and fecundity variation including the dynamics type of each colony as a random model.

