## Supplementary Information for "Population-level sensitivity to landscape variables reflects individual-based habitat selection in a woodland bat species"


Pierre-Loup Jan[1,2,*,†], Diane Zarzoso-Lacoste[3,4,5,*], Damien Fourcy[1], Alice Baudouin[3], Olivier Farcy[6], Josselin Boireau[7], Pascaline Le Gouar[3], Sébastien J. Puechmaille[8,9], Eric J. Petit[1]

[1] *ESE, Ecology and Ecosystem Health, INRAE, Agrocampus Ouest, 35042 Rennes cedex, France*
[2] *UMR 7372 CEBC, CNRS, La Rochelle Université, 79360 Villiers-en-Bois, France*
[3] *UMR 6553 ECOBIO, Université Rennes 1, Campus de Beaulieu, 35042 Rennes cedex, France*
[4] *UMR 8079 ESE, Université Paris-Sud/CNRS/AgroParisTech, Université Paris-Saclay, 91405 Orsay Cedex, France*
[5] *UMR 7058 EDYSAN, CNRS, Université de Picardie Jules Verne, 80037 Amiens Cedex 1, France*
[6] *Bretagne Vivante, 29221 Brest cedex 2, France*
[7] *Groupe Mammalogique Breton, 29450 Sizun, France*
[8] *Zoological Institute and Museum, Greifswald University, 17489 Greifswald, Germany*
[9] *ISEM, Univ Montpellier, CNRS, EPHE, IRD, Montpellier, France.*

* Co-first-authors

**Supplementary Information A:** Selection of population growth model

Growth rates can be estimated through different growth models, which can consider that the observed data are only the result of growth rate and observation error or can also account for other parameters, such as density dependence or carrying capacity. To perform model averaging in our dataset, we needed to determine which growth model would be the most appropriate to apply on all of our colonies.

For this purpose, we used the package "PVAclone" (Nadeem and Solymos 2016), which allows us to fit various population growth models to an empirical dataset. We tested six models commonly used and tested to describe population growth (Brook and Bradshaw 2006, Chamaillé-Jammes et al. 2008, Mech and Fieberg 2015): A Malthusian model, a Ricker model, a theta-logistic model, a logistic model, a Gompertz model, and a generalized Beverton-Holt model. The complete description of these models can be found in Nadeem and Solymos (2016), the Malthusian model being equivalent to a density-independent Ricker model (b parameter fixed to 0) and the logistic model being the theta-logistic model with the value of theta fixed to 1. We applied those six models on our whole dataset considering 5 clones for each colony, 20 000 iterations, and observation errors were assumed to follow a Poisson distribution. Models were then compared based on the proportion of colonies where the Markov chains of the models converged and the proportion of colonies where the models met were among the best fit for the data. We considered that models converged if the potential scale reduction factor was less than 1.1 (Brooks and Gelman 1998) and that a model was among the best fit for a given colony if its AICc was lower than the lowest AICc + 2 (Burnham and Anderson 2003).

**Table a: Comparison of growth model convergence and fit to observed count data in 94 colonies of *Rhinolophus hipposideros*. % convergence: Proportions of colonies with potential scale reduction factor < 1.1, % best fit : Proportions of colonies with ΔAICc < 2**

| Growth model | % convergence | % best fit |
|---|---|---|
| Malthusian model | 100% | 53.1% |
| Ricker model | 88.3% | 22.3% |
| Theta-logistic model | 68.1% | 21.3% |
| Logistic model | 86.2% | 37.2% |
| Gompertz model | 76.6% | 0% |
| Beverton-Holt model | 85.1% | 8.5% |

While the fit of the model seems to be highly colony dependent, these results show that the density-independent model is the only model that converged for the whole dataset and was considered to be among the best model for more than half of the colonies. Our dataset is composed of 94 colonies of different sizes, different observation durations, and most likely includes colonies at equilibrium as well as colonies that have not reached it at the end of the observation. This could explain why the models with best fit can be highly variable between colonies. The density-independent model is the simplest one, having a growth rate as the only parameter, and does not allow us to consider parameters such as density dependence or carrying capacity. However, the convergence and fitting results show that investigating the effects of environmental covariates at this scale would be more appropriate by considering the most parsimonious model. Subsequent analysis also accentuated the relevance of the density independent model because the model averaging performed on the growth rate provided by this model provided much more repeatable results than for any other parameter of the other models (data not shown).

**<u>Supplementary Information B</u>:** Aggregation of 38 cover types in six-landscape class.

| Cover type | Landscape class |
|---|---|
| Broadleaved trees seedling forest<br>Mixed seedling forest<br>Broadleaved trees seedling forest and coppice<br>Coppice<br>Open woodland<br>Poplar grove<br>Wooded area and hedgerows<br>Orchards<br>Undefined woodland | **Broadleaved woodland** |
| Conifers seedling forest<br>Coniferous seedling forest and coppice | **Coniferous woodland** |
| Building<br>Road<br>Urbanized areas<br>Industrial areas<br>Mine and dump<br>Public green area | **Artificial** |
| Undefined crop land<br>Cereal crop<br>Rapeseed crop<br>Various crop<br>Oilseed crop<br>Seeds<br>Vines<br>Arable land<br>Permanent crop | **Crop land** |
| Grassland<br>Heterogeneous agricultural area<br>Herbaceous and shrub vegetation area<br>Open area<br>Inland Wetland<br>Coastal Wetland<br>Heterogeneous area<br>Coast<br>Meadows and pastures<br>Set-aside land<br>Heathland | **Open land** |
| Freshwater | **Water bodies** |

**<u>Supplementary Information C</u>:** Description of the complete models used for model averaging.

Climatic variables were included in the model to account for their effect on colony size and fecundity (Jan et al. 2017) but were not tested through model averaging, and are presented in italics. * : Growth rate values were weighted according to the inverse of the standard deviation given by their estimates.

**Colony size complete model:**

Colony_size ~ %LAND_Broadleaved_woodland + %LAND_Coniferous_woodland + %LAND_Artificial + %LAND_Crop_land + %LAND_Open_land + %LAND_Water_bodies + MSI_Broadleaved_woodland + MSI_Coniferous_woodland +MSI_Artificial + MSI_Crop_land + MSI_Open_land + MSI_Water_bodies + SHEI + Mean_Patch_Size + *June_Precipitation + October_Precipitation + May_Temperature + November_Temperature*.

**Fecundity complete model:**

Fecundity ~ %LAND_Broadleaved_woodland + %LAND_Coniferous_woodland + %LAND_Artificial + %LAND_Crop_land + %LAND_Open_land + %LAND_Water_bodies + MSI_Broadleaved_woodland + MSI_Coniferous_woodland +MSI_Artificial + MSI_Crop_land + MSI_Open_land + MSI_Water_bodies + SHEI + Mean_Patch_Size + *April_Precipitation + October_Precipitation + April_Minimum_Temperature + July_Minimum_Temperature*.

**Growth rate complete model:**

Growth_rate* ~ %LAND_Broadleaved_woodland + %LAND_Coniferous_woodland + %LAND_Artificial + %LAND_Crop_land + %LAND_Open_land + %LAND_Water_bodies + MSI_Broadleaved_woodland + MSI_Coniferous_woodland +MSI_Artificial + MSI_Crop_land + MSI_Open_land + MSI_Water_bodies + SHEI + Mean_Patch_Size

**Supplementary Information D:** Effect size plots for the three best models investigating the effect of landscape on colony size (A), fecundity (B), and growth rate (C) in *Rhinolophus hipposideros* maternity colonies. Effect size were plotted for each model with the help of the R package *effects* (Fox 2003). Effect sizes for the growth rate analysis come from the model with the lowest AICc out of the ten runs. Colony size is represented on a logarithmic scale, and all values of landscape covariates (every X-axis) are scaled.

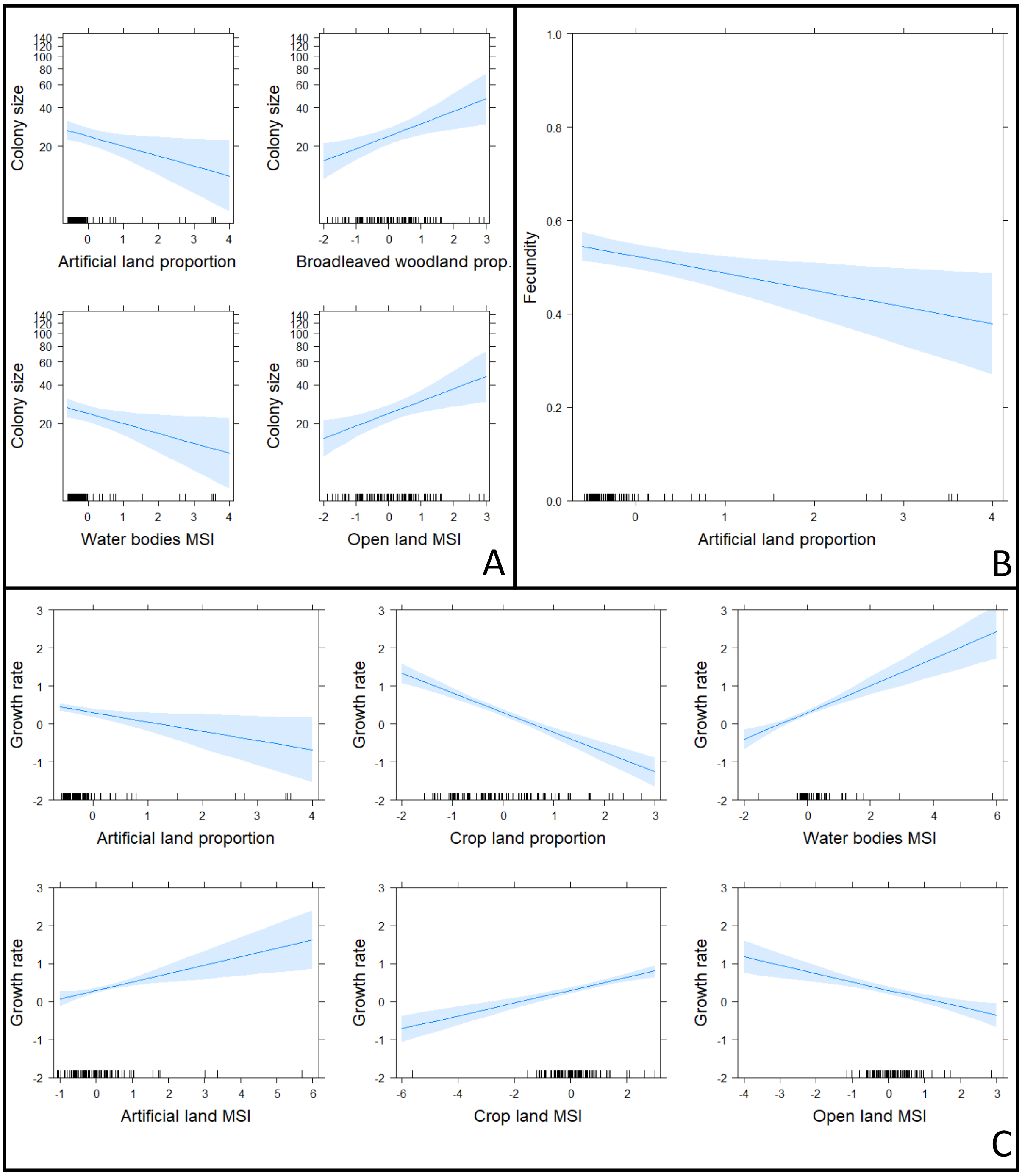

**<u>Supplementary Information E</u>:** Analysis of colony size and fecundity variation including the dynamics type of each colony as a random model.

To test if the dynamic pattern of our maternity colonies during the monitoring period would influence our result on colony size or fecundity, we assigned each colony to one of three categories, based on the fitting of Malthusian model, the fitting of logistic model, and the standard error around the growth rate. Those three categories are :

Stable colonies: (Lower AICc for Malthusian model than the logistic one, growth rate not significantly different from 0) – 52 colonies in this category.

Colonies with Malthusian growth: (Lower AICc for Malthusian model than the logistic one, growth rate significantly different from 0) – 11 colonies in this category.

Colonies with logistic growth: (Lower AICc for logistic model than the Malthusian one) – 31 colonies in this category.

We then replaced the (generalized) linear models used in our model averaging analysis by the corresponding mixed model, including the assignment to each category as a random effect. Results of this analysis is presented in the following tables.

**Table E1:** Results of a model averaging of population dynamics parameters as a function of the proportion of land (%LAND) and Mean Shape Index (MSI) of different cover type, the mean patch size and SHEI (Shannon's evenness index). Models also included the "dynamics type" as a random effect (see above). NS: non-significant predictor; '+': positive significant predictor; '-': negative significant predictor. When results differ between the two buffer sizes, results given by the 2500m buffer are indicated between brackets. Cells with significant results with the 500m buffer are shaded.

| | Broadleaved woodland | | Coniferous woodland | | Artificial | | Crop land | | Open land | | Water bodies | | SHEI | Mean patch size |
|---|---|---|---|---|---|---|---|---|---|---|---|---|---|---|
| | %LAND | MSI | %LAND | MSI | %LAND | MSI | %LAND | MSI | %LAND | MSI | %LAND | MSI | | |
| Colony size | + | NS | NS | NS | - (NS) | NS | NS | NS | NS | - (NS) | NS | + (NS) | NS | NS |
| Fecundity | NS | NS | NS | NS | - (NS) | NS | NS | NS | NS | NS | NS | NS | NS | NS |

**Table E2**: Coefficients, Residual degree of freedom (df), AICc and $R^2$ of the models investigating Colony size and fecundity variation, but with the type of dynamics included as random effect (see above). $R^2$ of models that include significant climatic predictor detected in a previous study (Jan et al. 2017) are also presented.

| Response variable | Radius | Fixed effects (Coefficients) | df | AICc | $R^2$ | $R^2$ with climatic variables |
|---|---|---|---|---|---|---|
| Colony size | 500 m. | Broadleaved woodland (0.23) and artificial (-0.18) proportion<br>Open land (-0.24) and water bodies (0.20) MSI | 87 | 777.4 | 0.32 | 0.35 |
| | 2500 m. | Broadleaved woodland (0.17) proportion | 92 | 798.7 | 0.003 | 0.04 |
| Fecundity | 500 m. | Artificial (-0.04) proportion | 90 | -114.47 | 0.08 | 0.14 |
| | 2500 m. | - | - | - | - | - |
